# Strain-resolved metagenomics shows gut microbiome signatures in a case-control study of Malawian children with zoonotic schistosomiasis

**DOI:** 10.64898/2026.11.25.769124

**Authors:** Angus M. O’Ferrall, David Lally, Peter Makaula, Gladys Namacha, Lucas J. Cunningham, John Archer, Joseph M. Lewis, Andrew Bennett, Simon C. Wagstaff, Sekeleghe A. Kayuni, Janelisa Musaya, J. Russell Stothard, Adam P. Roberts

**Affiliations:** Department of Tropical Disease Biology, Liverpool School of Tropical Medicine, Liverpool, United Kingdom; Malawi Liverpool Wellcome Research Programme, Queen Elizabeth Central Hospital, Blantyre, Malawi; Department of Clinical Sciences, Liverpool School of Tropical Medicine, Liverpool, United Kingdom; Research Computing Unit, Liverpool School of Tropical Medicine, Liverpool, United Kingdom; Community and Environmental Health Department, School of Global and Public Health, Kamuzu University of Health Sciences, Blantyre, Malawi; Department of Pathology, School of Medicine and Oral Health, Kamuzu University of Health Sciences, Blantyre, Malawi

**Author notes:** Corresponding author –.

## Abstract

In 2024, Mangochi District, Malawi, we identified unexpected cases of zoonotic schistosomiasis associated with faecal egg excretion of *Schistosoma haematobium*, *Schistosoma mattheei* and hybrids thereof (*S. haematobium* × *S. mattheei*). Schistosomiasis is associated with gut microbiome alterations, but these associations remain unexplored in zoonotic schistosomiasis. Whether people with schistosomiasis are colonised with more closely related gut bacterial strains than people without schistosomiasis is also unclear. To better understand these patterns, we conducted a case-control study, using strain-resolved metagenomics to analyse faecal samples from 36 children: 12 with zoonotic schistosomiasis (*S. haematobium* and *S. mattheei*), 12 with ‘classic’ urogenital schistosomiasis (*S. haematobium* only) and 12 uninfected controls. Microbial diversity, taxonomic composition, metabolic pathways and antimicrobial resistance gene profiles did not differ significantly between groups. However, *Brachyspira*, particularly *Brachyspira aalborgi* – an organism often associated with intestinal spirochaetosis – was more abundant in children with zoonotic schistosomiasis. We found no evidence of closely related *B. aalborgi* strains shared between participants, suggesting that localised *B. aalborgi* transmission networks are unlikely to explain the enrichment. In contrast, phylogenetic profiling across 143 other bacterial species revealed greater strain similarity between the gut microbiomes of children with any type of schistosomiasis than between those of controls. Broadly, we demonstrate the scientific value of strain-resolved metagenomics for investigating linked bacterial and schistosome exposures where local water, sanitation and hygiene infrastructure is inadequate. Future inclusion of livestock and environmental samples could be used to better identify where transmission pathways overlap, providing evidence to inform integrated disease surveillance and control.

**Significance statement:** Zoonotic schistosome transmission is geographically localised, occurring where humans, animals and freshwater snails share freshwater environments and where water, sanitation and sanitation (WASH) infrastructure is lacking. These factors may also facilitate exposure to enteric bacteria, but potential links between parasitic infection and the distribution of gut bacterial strains remain poorly understood. Using strain-resolved metagenomics, we identified enrichment of intestinal spirochaetes associated with zoonotic *Schistosoma mattheei* infection and greater gut bacterial strain similarity among children with schistosomiasis. These exploratory findings highlight how studying bacterial strains, rather than microbiome composition alone, can reveal epidemiological patterns potentially reflecting shared environmental exposures. This approach could help inform integrated strategies to reduce exposure to parasitic and bacterial pathogens in communities with inadequate WASH infrastructure.

## Introduction

Across much of sub-Saharan Africa (SSA), reliance on unsafe water for essential daily processes, such as drinking and washing, predisposes underserved populations to several waterborne diseases. Schistosomiasis, for example, accounts for an estimated loss of 1.8 million disability-adjusted life-years annually, with the majority of this burden occurring in SSA, where it remains one of the region’s most pervasive parasitic diseases [1]. The disease is caused by infection with blood flukes of the genus *Schistosoma* and is transmitted via contact with infectious larvae released from freshwater snail intermediate hosts [1]. Mature *Schistosoma* worms are considered to be relatively benign within the vasculature of human hosts, but their eggs can become trapped in host tissues, where they induce inflammation and tissue damage [1]. Host immune and inflammatory responses, and egg-induced micro-perforations, are thought to contribute to gut microbiome alterations [2]. This may be clinically relevant, as gut inflammation can promote enrichment of opportunistic bacterial pathogens, including those associated with a substantial burden of antimicrobial resistance (AMR) [3,4]. Furthermore, the gut microbiome can influence host immune responses to infection and the metabolism and efficacy of orally administered drugs, potentially including praziquantel, the principal treatment for schistosomiasis [5–7].

There are two major clinical manifestations of schistosomiasis. Intestinal schistosomiasis is predominantly caused by *Schistosoma mansoni* in Africa and the Americas, and by *Schistosoma japonicum* in Asia, whereas *Schistosoma haematobium* is the principal cause of urogenital schistosomiasis, which is particularly prevalent in SSA [1]. Despite not typically being thought to cause significant intestinal disease, *S. haematobium* infection has been associated with gut microbiome alterations [8]. Notably, relatively few differentially abundant microbial taxa have been consistently identified across gut microbiome studies of schistosomiasis [2]. This lack of concordance may reflect intravascular localisation patterns of the different *Schistosoma* species, involving either the mesenteric venous system (intestinal schistosomiasis) or the vesical venous plexus (urogenital schistosomiasis), leading to differences in the predominant sites of egg deposition. The immunopathology associated with tissue-trapped eggs in these distinct anatomical sites may create different systemic or organ-specific inflammatory responses [1], potentially contributing to species-specific microbiome signatures. However, modest sample sizes, geographic variation in diet, nutrition and co-infection patterns, as well as differences in sequencing and analytical methods, are all likely to contribute to the heterogeneity of reported findings.

*Schistosoma mattheei* is a zoonotic species within the *S. haematobium* group that most commonly infects livestock, causing intestinal schistosomiasis [9]. However, it is occasionally reported to cause urogenital schistosomiasis in humans, as eggs of *S. mattheei* are more readily detected in urine than in faecal samples. In 2024, we reported on cases of schistosomiasis associated with the unexpected faecal egg excretion of *S. haematobium*, *S. mattheei* and hybrids thereof (*S. haematobium* × *S. mattheei*) in people from Samama, Mangochi District, Malawi [10]. Urogenital schistosomiasis caused by *S. haematobium* is highly endemic in this community, and although *S. mansoni* infections have also been reported over the last decade, typical cases of intestinal schistosomiasis are decreasing in prevalence [11–13]. Nonetheless, we subsequently deployed novel DNA-based molecular diagnostics to reveal that the detection of *S. haematobium* mitochondrial DNA (mtDNA) was more common than the detection of *S. mansoni* mtDNA in the faeces of school-aged children (SAC) in Samama, and was significantly associated with *S. mattheei* co-infection [14]. These findings highlighted complex co-infection dynamics that may influence schistosome tissue tropism, egg excretion routes and potentially gut microbiome profiles. Furthermore, localised patterns of enteric bacterial transmission in Samama mirror the focal epidemiology of zoonotic schistosome infections and indicate that shared environmental exposures may contribute to the distribution of both parasites and enteric bacteria [15–17].

Here, we report the findings of a case-control study comparing the gut microbiomes of SAC with zoonotic schistosomiasis and ‘classic’ urogenital schistosomiasis with those from uninfected controls in the same village from a subset of faecal DNA extracts recently used to investigate the molecular epidemiology of intestinal schistosomiasis in Samama [14]. We leverage the high resolution of shotgun metagenomic sequencing data to demonstrate the utility of strain-resolved analyses that provide insights into potentially linked pathways of bacterial and schistosome exposure in settings with inadequate water, sanitation and hygiene (WASH) infrastructure.

## Results

### Participant characteristics

The zoonotic schistosomiasis (‘ZS’) group comprised 12 SAC (5 females and 7 males), from whom *Schistosoma* spp. ribosomal internal transcribed spacer 2 (ITS2) DNA was detected in the faeces by real-time PCR, and both *S. haematobium* and *S. mattheei* mtDNA signatures, but not *S. mansoni*, were detected in the faeces and/or urine by high-resolution melt (HRM) real-time PCR. Nine of these SAC excreted detectable *Schistosoma* eggs in the urine that were identified by urine filtration and microscopy, while co-infection with *Giardia duodenalis* (syn. *Giardia intestinalis*, *Giardia lamblia*), another common endemic waterborne parasite, was detected in three faecal samples from SAC in the ZS group by real-time PCR. The 12 SAC in the ZS group were matched by sex, age and *G. duodenalis* infection status to two groups of 12 SAC from the same village: an uninfected ‘Control’ group, and a ‘classic’ urogenital schistosomiasis (‘UGS’) group, from whom *S. haematobium* eggs and DNA were detected on urine filters but no *Schistosoma* DNA was detected in faeces. Participants from the three groups were dispersed across the village, with no obvious geographic clustering (*Supplementary Figure 1*). No differences in antibiotic usage in the six months prior to sample provision, use of piped water as the primary drinking water source, daily contact with environmental water bodies (including river, lake, pond, marsh, swamp and rice paddy water) or latrine use for defaecation were detected between groups (*Table 1*).

**Table 1.** Participant characteristics at the time of sample collection. Age (in years) is presented as median (IQR). Categorical variables are presented as proportions (%). *P*-values were calculated to assess differences in participant characteristics between groups using a Kruskal–Wallis test for age (numerical variable) and two-sided Fisher’s exact tests for categorical variables.

| Variable | Control<br>(n = 12) | UGS<br>(n = 12) | ZS<br>(n = 12) | *P*-value | Total<br>(n = 36) |
|---|---|---|---|---|---|
| Age (years) | 10<br>(7.00–11.50) | 11<br>(6.75–12.25) | 10<br>(7.00–11.50) | 0.991 | 10<br>(7.00–12.25) |
| Sex: female | 41.7<br>(5/12) | 41.7<br>(5/12) | 41.7<br>(5/12) | 1.000 | 41.7<br>(15/36) |
| Giardiasis positive | 25.0<br>(3/12) | 25.0<br>(3/12) | 25.0<br>(3/12) | 1.000 | 25.0<br>(9/36) |
| Antibiotic use within six months | 41.7<br>(5/12) | 50.0<br>(6/12) | 25.0<br>(3/12) | 0.576 | 38.9<br>(14/36) |
| Primary drinking water source: piped water | 50.0<br>(6/12) | 58.3<br>(7/12) | 25.0<br>(3/12) | 0.334 | 44.4<br>(16/36) |
| Daily environmental water contact | 91.7<br>(11/12) | 91.7<br>(11/12) | 91.7<br>(11/12) | 1.000 | 91.7<br>(33/36) |
| Latrine use for defaecation | 91.7<br>(11/12) | 83.3<br>(10/12) | 83.3<br>(10/12) | 1.000 | 86.1<br>(31/36) |

### Gut microbiome community composition and diversity are similar across schistosomiasis categories

Faecal DNA underwent shotgun metagenomic sequencing; the number of 150 bp Illumina read pairs per sample ranged from 41,125,582 to 59,224,576 (*Supplementary Table 1*). In total, 19 bacterial phyla were detected across all faecal DNA samples (*Figure 1a*). Of these, the most abundant was Firmicutes (mean relative abundance = 50.9%), followed by Bacteroidota (mean relative abundance = 34.1%). Pseudomonadota (syn. Proteobacteria) was the third most abundant phylum, but with much lower levels than the two dominant phyla (mean relative abundance of 6.2%). At lower taxonomic levels, 354 orders, 1,016 genera and 1,575 species of bacteria were detected across samples. Kruskal–Wallis tests did not detect any differences in alpha diversity metrics between groups (Shannon: *P* = 0.479; Gini–Simpson: *P* = 0.145; Richness: *P* = 0.703) (*Figure 1b*).

**Figure 1.**
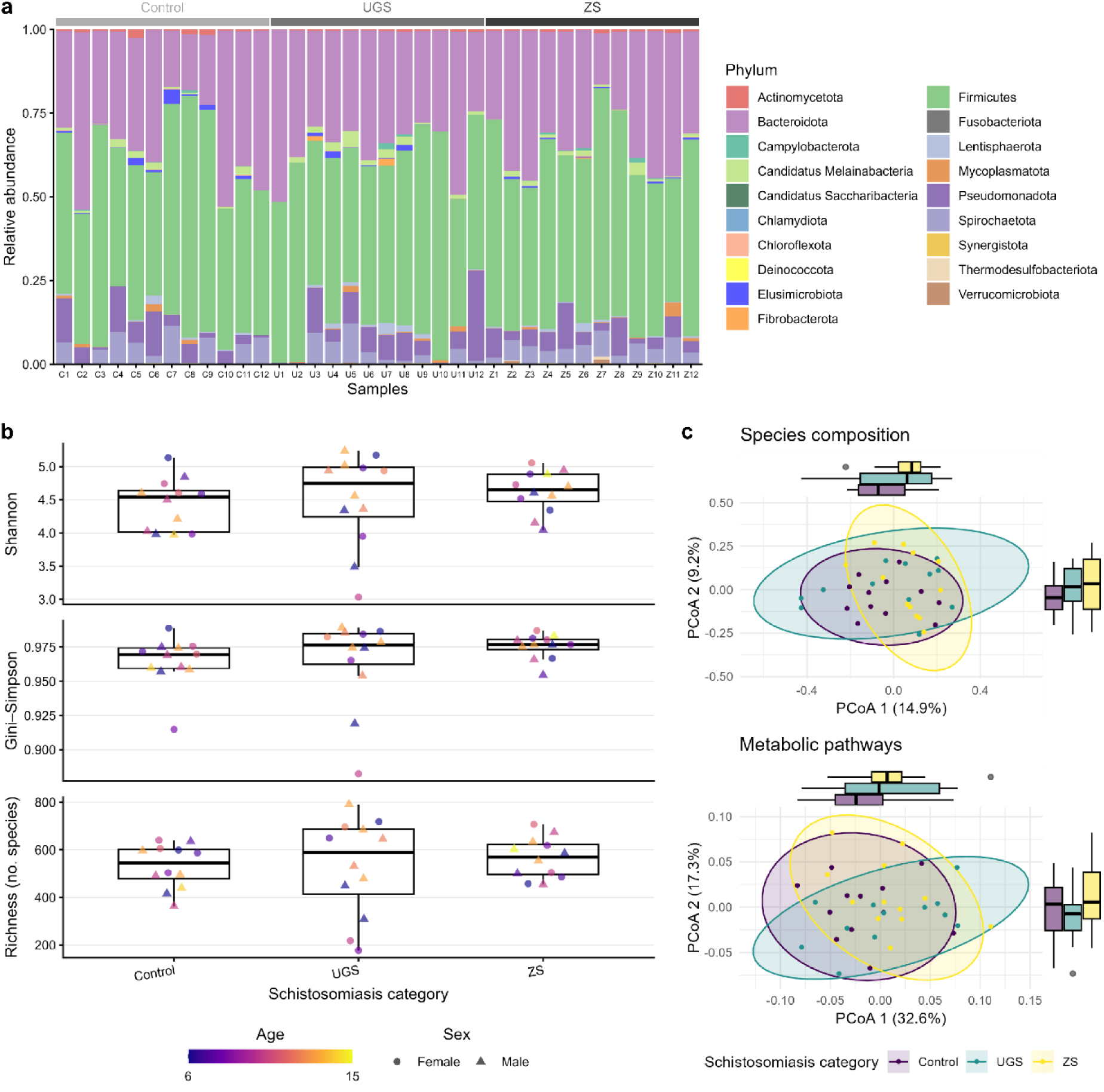
Gut microbiome composition and diversity comparisons between schistosomiasis categories. (a) Total sum-scale (TSS)-normalised relative abundance of bacterial phyla detected in faecal samples using MetaPhlAn. Samples from the ‘Control’, ‘UGS’ and ‘ZS’ groups begin with ‘C’, ‘U’ and ‘Z’, respectively. (b) Alpha diversity statistics by schistosomiasis category. Point colour indicates participant age and shape indicates participant sex. Kruskal–Wallis tests did not detect any differences in alpha diversity metrics between groups. (c) Principal Coordinate Analysis (PCoA) plot of all-versus-all Bray–Curtis dissimilarity (beta diversity) for species composition and metabolic pathways, with 95% multivariate t-distribution ellipses stratified by schistosomiasis category. Values in axis parentheses indicate the percentage of overall variance in the data that is captured by the axis.

Principal Coordinate Analysis (PCoA) of species-level taxonomic composition and metabolic pathway profiles did not show any distinct clustering of samples by schistosomiasis category (*Figure 1c*). Associations of sample metadata with microbiome community composition and function were further assessed with Permutational Multivariate Analysis of Variance (PERMANOVA) testing upon Bray–Curtis dissimilarity matrices of species- and genus-level taxonomic profiles, and metabolic pathways. No variables were significantly associated with species composition. Only age group, where participants were split into categories of 6–10 and 11–15 years, was significantly associated with genus composition (R^2^ = 0.046, *P* = 0.040) and metabolic pathway profiles (R^2^ = 0.073, *P* = 0.010), while schistosomiasis category, sex, giardiasis, antibiotic use within the previous six months and use of piped water as the primary drinking water source were not statistically associated with either taxonomic or functional profiles (*Table 2*). Homogeneity of multivariate dispersion was confirmed for the significant age group comparisons (genus composition: *P* = 0.160; metabolic pathways: *P* = 0.969).

**Table 2.** Permutational Multivariate Analysis of Variance (PERMANOVA) examining associations between participant characteristics and gut microbiome bacterial species composition, genus composition and metabolic pathway profiles. Covariates included in the analysis were selected based on potential biological relevance and sufficient variability across participants. *Statistical significance was defined as *P* <0.05.

| Variable | Species composition |  | Genus composition |  | Metabolic pathways |  |
|---|---|---|---|---|---|---|
|  | R<sup>2</sup> | *P*-value | R<sup>2</sup> | *P*-value | R<sup>2</sup> | *P*-value |
| Schistosomiasis category | 0.048 | 0.812 | 0.047 | 0.685 | 0.044 | 0.627 |
| Age group<br>(6–10 or 11–15) | 0.038 | 0.084 | 0.046 | 0.040* | 0.073 | 0.010* |
| Sex | 0.036 | 0.118 | 0.045 | 0.050 | 0.044 | 0.110 |
| Giardiasis | 0.021 | 0.892 | 0.020 | 0.817 | 0.011 | 0.930 |
| Antibiotic use within six months | 0.031 | 0.286 | 0.027 | 0.451 | 0.031 | 0.293 |
| Primary drinking water source (piped or non-piped) | 0.031 | 0.287 | 0.034 | 0.197 | 0.044 | 0.105 |

### Tetracycline resistance genes dominate the gut resistome

Reads mapped with >90% nucleotide identity and >80% template coverage to 112 ARG references in the ResFinder database. These ARG references belonged to 68 Cluster90s (defined as groups of highly similar ARG reference sequences clustered at a 90% nucleotide identity threshold), predicted to confer resistance to 15 antibiotic classes (*Supplementary Figure 2*). Tetracycline resistance genes contributed the greatest ARG load across all groups (*Figure 2a*). Tetracycline resistance genes were estimated to account for the largest proportion of the resistome in all samples by class, with a mean relative abundance of 57.34% across all samples, followed by beta-lactam and macrolide-lincosamide-streptogramin B (MLSB) resistance genes, with a mean relative abundance of 24.09% and 15.55%, respectively. Assemblies recovered 51 of the 112 ARGs that were identified by read-mapping (*Figure 2b*), 18 of which could be binned to metagenome-assembled genomes (MAGs) (*Supplementary Figure 3*). No extended-spectrum beta-lactamase (ESBL) genes were detected either by read-mapping or assembly-based methods, despite the carriage of *bla*_CTX-M_-harbouring *E. coli* being highly prevalent in this setting at the time of sampling (around 50%), as demonstrated in our recent selective culture-based whole-genome sequencing study [15]. No ARG classes or Cluster90s were differentially abundant in the ZS or UGS groups compared to the Control group using a permissive exploratory threshold of *q* <0.10.

**Figure 2.**
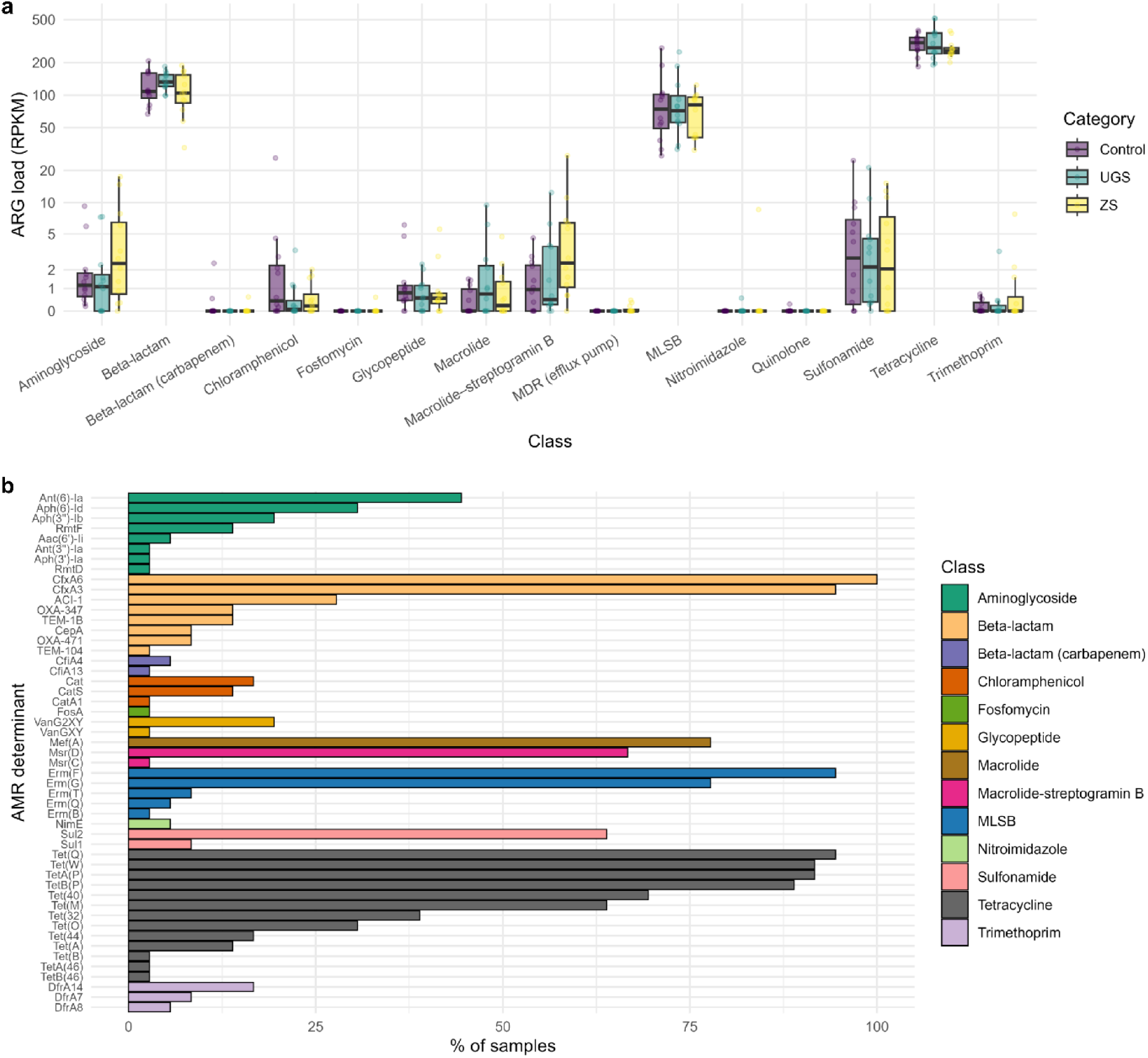
Antimicrobial resistance (AMR) determinants identified in the gut microbiota. (a) Antimicrobial resistance gene (ARG) load of detected ARGs predicted to confer resistance to each antibiotic class, stratified by schistosomiasis category. No ARG classes were associated with either zoonotic schistosomiasis or urogenital schistosomiasis. (b) Prevalence of AMR determinants identified in metagenome assemblies, coloured by antibiotic class. MDR = multidrug-resistant; MLSB = macrolide-lincosamide-streptogramin B; RPKM = Reads Per Kilobase per Million mapped bacterial reads.

### Intestinal spirochaetes are enriched in zoonotic schistosomiasis

Differential abundance analysis revealed enrichment of bacteria from the *Brachyspira* genus in the gut microbiomes of participants from the ZS group compared with controls. This association was detected using both our primary analytical method, MaAsLin2 (Microbiome Multivariable Associations with Linear Models) [18] (log_2_ fold change [log2FC] relative abundance = 3.41, SE = 0.86, *P* <0.001, *q* = 0.045), and in a sensitivity analysis using LinDA (Linear Models for Differential Abundance Analysis) [19] (log2FC abundance = 3.68, SE = 0.76, *P* <0.001, *q* = 0.003). No phyla, other genera or metabolic pathways met either the *q* <0.05 significance threshold or a more permissive exploratory threshold of *q* <0.10 with either differential abundance analysis method for the ZS or UGS groups compared to the Control group.

Two *Brachyspira* species were identified by MetaPhlAn profiling: *Brachyspira aalborgi* (3/12 Control, 6/12 UGS and 8/12 ZS) and *Brachyspira pilosicoli* (0/12 Control, 1/12 UGS and 5/12 ZS). Co-colonisation of *B. aalborgi* and *B. pilosicoli* was identified in four participants (0/12 Control, 1/12 UGS and 3/12 ZS). *Brachyspira aalborgi* was the more abundant of the two species and appeared to be the main driver of the differential abundance signal. However, among participants in the ZS group, no significant correlation was identified between normalised *Schistosoma* DNA abundance and *Brachyspira* spp., *B. aalborgi* or *B. pilosicoli* relative abundance in faeces (*Figure 3a*). StrainPhlAn reconstructed a marker-gene phylogeny of the dominant *B. aalborgi* strain from each sample in which the species was detected, with the exception of one sample from the UGS group that had insufficient marker-gene coverage. The reconstructed dominant *B. aalborgi* strains were phylogenetically distinct between participants (*Figure 3a*). Owing to the low relative abundance (<0.1%) of *B. pilosicoli* across all but one sample, marker gene coverage was insufficient to perform StrainPhlAn phylogenetic reconstruction.

**Figure 3.**
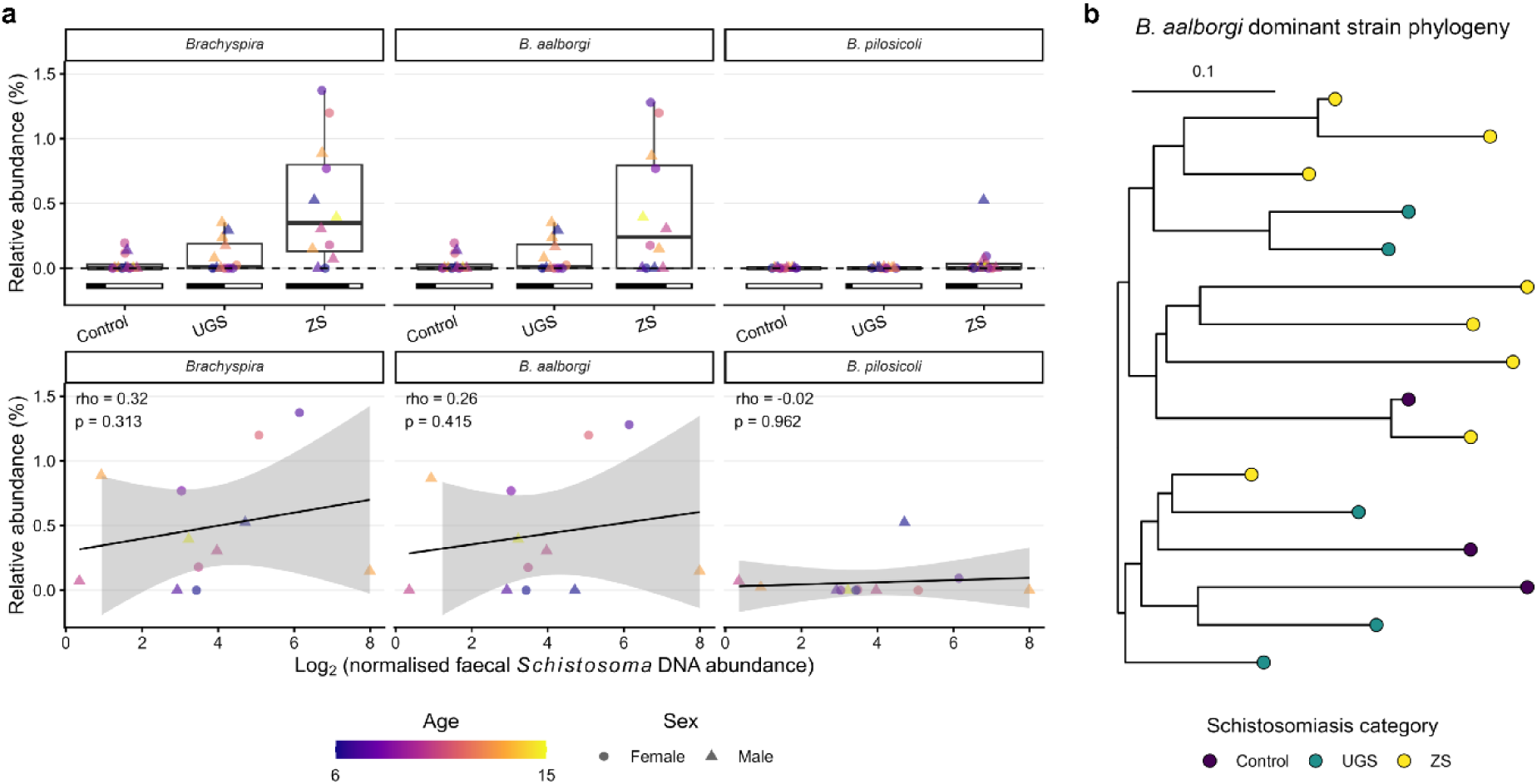
Associations between *Brachyspira* colonisation and zoonotic schistosomiasis. (a) Relative abundance of the genus *Brachyspira* and the species *Brachyspira aalborgi* and *Brachyspira pilosicoli*, stratified by schistosomiasis category (top), and their relationship with normalised faecal *Schistosoma* DNA abundance among children with zoonotic schistosomiasis (bottom). MaAsLin2 models estimated a log_2_ fold change (log2FC) in relative abundance of 3.41 in zoonotic schistosomiasis compared to controls (*P* <0.001, *q* = 0.045). Boxplots show the median and IQR, whiskers extend to 1.5 × IQR, and points represent individual participants. In the scatterplots, lines represent linear regression fitted to log₂-transformed normalised faecal *Schistosoma* DNA abundance, with shaded areas indicating the 95% confidence interval. Spearman’s rank correlation coefficients (rho) and corresponding *P*-values are shown. Point colour indicates participant age and point shape indicates sex. (b) Maximum-likelihood phylogeny of dominant *B. aalborgi* strains reconstructed using StrainPhlAn. Tip colours indicate schistosomiasis category. The scale bar indicates substitutions per site across species-specific marker genes.

Six high-quality MAGs from the genus *Brachyspira* were recovered across samples (five *B. aalborgi* and one *B. pilosicoli*), each belonging to a distinct subspecies group (all pairwise average nucleotide identity [ANI] estimates <98%). We mapped reads from all samples to these six references using TRACS (TRAnsmission Clustering of Strains) [20] to estimate the lower bound of the number of single nucleotide polymorphisms (SNPs) between strains colonising different participants. After masking potential recombination regions in alignments, the least divergent pair of *B. aalborgi* strains differed by 4,625 SNPs across 2.14 Mbp of alignment. Therefore, no evidence for *B. aalborgi* transmission networks involving multiple study participants was identified. Insufficient coverage precluded equivalent analyses for *B. pilosicoli*, which was less abundant and prevalent than *B. aalborgi*.

### Children with schistosomiasis harbour more similar dominant bacterial strains

Of the 860 MetaPhlAn species-level genome bins (SGBs) with a prevalence >20% in this study, StrainPhlAn reconstructed marker-gene phylogenies of the dominant strains for 518 SGBs. Of these, 143 SGBs met the minimum requirements for estimation of SGB-specific genetic distance thresholds to infer sharing of highly similar strains. These SGBs were retained for downstream strain-level similarity analyses. Between all possible pairwise combinations of participants from different households (n = 627), the median number of SGBs identified in both participants was 54 (IQR = 47–64), from the 143 well-represented SGBs included.

Median pairwise strain-level microbiome similarity (the percentage of SGBs profiled in both samples in which dominant strains were deemed highly similar by StrainPhlAn) was 9.38% (IQR = 6.06–12.96). Kruskal–Wallis tests indicated that strain-level microbiome similarity differed according to the schistosomiasis infection status of participant pairs (both infected, neither infected or discordant: *P* = 0.001), but not according to schistosomiasis infection type (both UGS, both ZS or discordant: *P* = 0.092). Pairs of participants from different households in which both individuals had schistosomiasis shared more similar strains than discordant pairs and pairs in which neither participant was infected (*Figure 4*). In a participant-level sensitivity analysis, participants with schistosomiasis exhibited a greater within-group relative to between-group strain-level microbiome similarity than participants without schistosomiasis. The median within-minus-between difference in strain-level microbiome similarity was 1.82 percentage points among participants with schistosomiasis, compared with −0.62 percentage points among uninfected participants. The difference between groups was statistically significant (Wilcoxon rank-sum test: *P* = 0.003).

**Figure 4.**
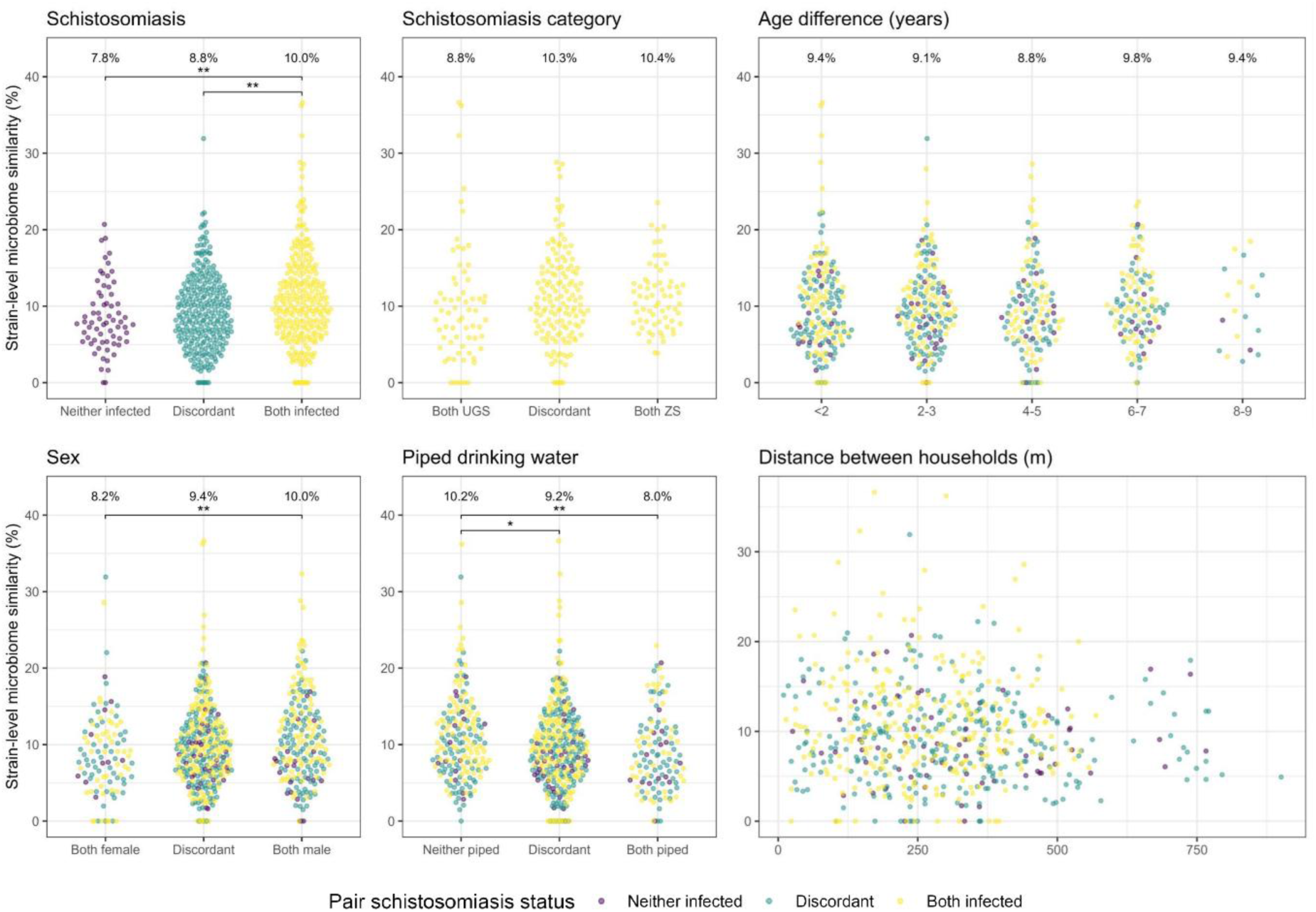
Associations between pairwise strain-level microbiome similarity and pair characteristics. Pairwise strain-level microbiome similarity was calculated as the percentage of species-level genome bins (SGBs) in which the dominant strains were classified as highly similar by StrainPhlAn. Pairwise comparisons were made for the following variables: schistosomiasis, schistosomiasis category (among participants with schistosomiasis), age difference (years), sex, use of piped water as the primary drinking water source and geographic distance between households. Numbers above each category indicate the median strain-level microbiome similarity per group. Pairwise comparisons are coloured according to schistosomiasis infection status (neither infected, discordant or both infected). Statistical significance was assessed using Kruskal–Wallis tests followed by post-hoc Benjamini–Hochberg-adjusted Dunn’s tests for categorical variables and Spearman’s rank correlation for household distance. Statistically significant associations are shown by asterisks: *P* <0.05 (*), *P* <0.01 (**).

Additionally, primary drinking water source was associated with strain-level microbiome similarity (Kruskal–Wallis test: *P* = 0.006): pairs in which both participants primarily used non-piped drinking water were shown to share more similar strains. Participant-level sensitivity analyses supported this finding; participants primarily drinking non-piped water exhibited a greater within-group relative to between-group strain-level microbiome similarity, with a median within-minus-between difference of 1.05 percentage points, compared with −1.19 percentage points among participants primarily drinking piped water (Wilcoxon rank-sum test: *P* = 0.006). Sex was also associated with pairwise strain-level microbiome similarity (Kruskal–Wallis test: *P* = 0.010), with pairs of males exhibiting greater strain-level microbiome similarity than pairs of females. However, this association was not supported by participant-level sensitivity analyses, which found no significant difference in within-group relative to between-group strain-level microbiome similarity between males and females. No significant associations with pairwise strain-level microbiome similarity were observed for distance between households (Spearman’s rank correlation: rho = −0.06, *P* = 0.122), age difference category (Kruskal–Wallis test: *P* = 0.704) or giardiasis infection (Kruskal–Wallis test: *P* = 0.800).

The associations of schistosomiasis infection and primary drinking water source with pairwise strain-level microbiome similarity remained evident in a multivariable negative binomial regression model after adjustment for age difference, sex of pairs and distance between participant households (*Supplementary Table 2, Figure 5a*). The greatest positive incidence rate ratios (IRRs) were observed between pairs of people with schistosomiasis, regardless of whether the individuals in the pairs were in the UGS group or the ZS group. In the multivariable model, distance between households was negatively associated with strain-level microbiome similarity. Permutation-based variable importance analysis identified schistosomiasis status as the strongest contributor to model fit; breaking the relationship between this variable and the outcome produced the greatest reduction in model log-likelihood relative to the original model (*Figure 5b*).

**Figure 5.**
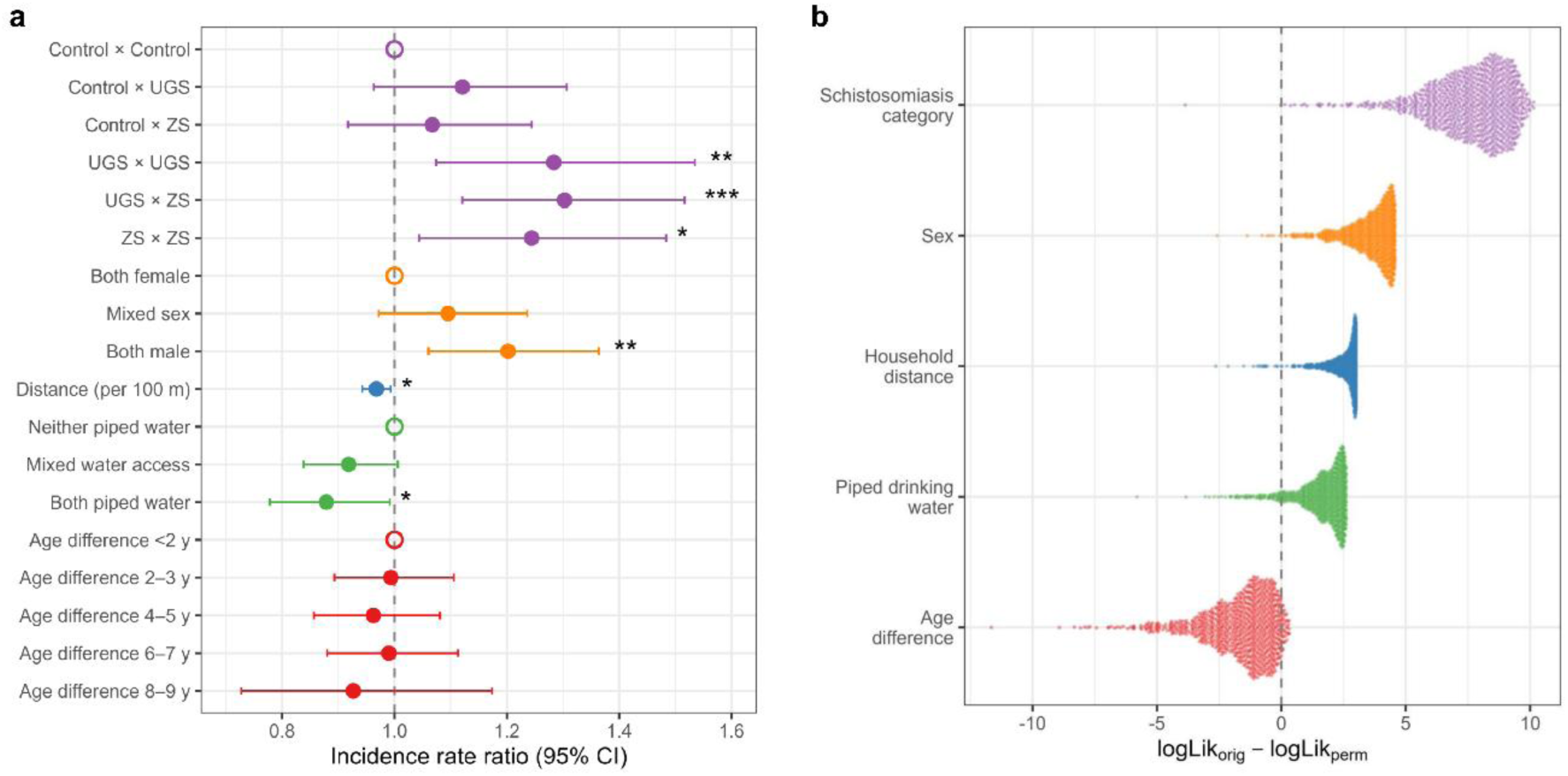
Multivariable negative binomial regression analysis of factors associated with strain-level microbiome similarity. (a) Incidence rate ratios (IRRs) and 95% confidence intervals from the multivariable negative binomial model investigating the number of highly similar dominant strains shared between participant pairs, with the logarithm of the total number of species-level genome bins (SGBs) identified in samples of both participants included as an offset. Reference categories are shown as open circles. Household distance is modelled per 100 m increase. Statistically significant associations are shown by asterisks: *P* <0.05 (*), *P* <0.01 (**), *P* <0.001 (***). (b) Permutation-based variable importance analysis showing the change in model log-likelihood after independently permuting each covariate across participant pairs (1,000 permutations per variable). Larger positive differences between the original and permuted model log-likelihoods indicate a greater contribution of that variable to model fit.

## Discussion

This study provides preliminary insights into associations between gut microbiome profiles and an emerging form of zoonotic schistosomiasis involving *S. mattheei* in southern Malawi. While overall gut microbiome diversity and community composition were similar between children with zoonotic schistosomiasis, ‘classic’ urogenital schistosomiasis and uninfected controls, an enrichment of *Brachyspira* bacteria was identified in the gut microbiomes of children with zoonotic schistosomiasis. Additionally, children with either form of schistosomiasis shared a greater proportion of similar dominant bacterial strains than uninfected individuals, suggesting that schistosomiasis may be associated with shared environmental exposures influencing bacterial strain distribution.

The observed microbiome alterations were predominantly taxon-specific, with no evidence of widespread dysbiosis, although the detection of subtle community-wide compositional shifts may only have been possible with a larger sample size. Nonetheless, the exploratory findings presented here are consistent with a recent gut microbiome study in Uganda, where typical intestinal schistosomiasis caused by *S. mansoni* in rural settings was not associated with significant beta diversity separation despite being associated with changes in the abundance of specific microbial taxa [21]. However, in that study, the altered taxa were proposed to influence host cardiovascular disease risk through microbe–metabolite networks, while we found no evidence of altered microbial metabolic pathways in *S. haematobium* group infections, which may be less associated with microbiome functional shifts. More broadly, published studies on the gut microbiome in schistosomiasis have produced inconsistent findings [2], likely reflecting differences in infecting *Schistosoma* species, infection intensity, participant ages, geography, diet, environmental exposures, sample size and the rapidly evolving analytical approaches used in microbiome research.

An important feature of this study population is the long-term participation of SAC in schistosomiasis surveillance and treatment programmes in Samama, Mangochi District, dating back to 2018 [11]. During the years preceding sample collection, participants underwent repeated praziquantel treatment and are therefore likely to have experienced multiple cycles of infection, treatment and reinfection with schistosomes. Consequently, the gut microbiome signatures observed here may reflect associations with relatively new *S. haematobium* group infections superimposed upon established gut microbiomes that are likely to have been influenced by various endemic parasite infections, including schistosomiasis and giardiasis, over a period of years. This may partially explain why we did not identify global shifts in bacterial community composition and function, contrasting with studies of younger children in which most participants had not previously received praziquantel treatment [8,22].

The analyses presented here did, however, identify an association between zoonotic schistosomiasis and enrichment of the genus *Brachyspira*, particularly *B. aalborgi*, in the gut microbiomes of affected children. *Brachyspira* spp. are oxygen-tolerant anaerobic spirochaetes that have been isolated from the gastrointestinal tracts of various mammals and birds [23]. Two *Brachyspira* species have been associated with human intestinal spirochaetosis, a cause of non-specific abdominal symptoms, including diarrhoea, abdominal pain and bloating [24]. Both of these species were detected here: *B. aalborgi* was predominant, whereas *B. pilosicoli* was identified at lower prevalence and abundance. Human intestinal spirochaetosis is characterised by colonisation of the colonic epithelium by *Brachyspira* spp. and is reported most frequently in immunocompromised individuals or alongside co-infections with other enteric pathogens, such as *Helicobacter pylori* [24,25]. However, detection of *Brachyspira* DNA in faecal samples alone does not confirm intestinal spirochaetosis, as diagnosis of invasive disease requires colonoscopy, biopsy and histopathological examination, while some cases of colonisation are asymptomatic [25]. As clinical assessments were not routinely performed in our study, we were unable to determine whether *Brachyspira* enrichment was associated with gastrointestinal symptoms or histological evidence of intestinal spirochaetosis.

Nevertheless, the enrichment of *Brachyspira* raises the possibility of a biological interaction between *S. haematobium* group infections, specifically *S. mattheei*, and colonisation by intestinal spirochaetes. One plausible explanation is that deposition of *Schistosoma* eggs around the intestines and the associated local inflammatory response alter the intestinal environment, facilitating colonisation and/or invasion with spirochaete bacteria. Alternatively, the presence of *Brachyspira* and other potentially immunomodulatory co-infections not tested for here may be linked with increased susceptibility to *S. mattheei* infection. This may include HIV, which is highly prevalent in Mangochi District alongside schistosomiasis [26], but was only accounted for by self-reporting in questionnaires in our study. Meanwhile, environments could simultaneously expose some people to both *S. mattheei* and *Brachyspira* spp., but the presence of distinct dominant *B. aalborgi* strains and the absence of evidence for recent transmission between participants in SNP-based analyses suggests that enrichment is less likely to reflect dissemination of specific strains at zoonotic schistosome transmission foci.

In contrast, strain-level analyses across common SGBs showed that children with schistosomiasis shared a greater proportion of similar dominant bacterial strains between each other than uninfected children, with schistosomiasis infection status emerging as the strongest predictor of strain-level microbiome similarity in multivariable analyses. Beyond conventional taxonomic analyses, these findings provide insights into shared epidemiological patterns that may link schistosome infection and the distribution of gut bacterial strains through common environmental exposures. A potential reason for the association is that children who recently acquired schistosomiasis are likely to have been exposed to environmental water bodies where contact with human and animal faecal contamination may also facilitate acquisition of similar enteric bacterial strains. Although questionnaire data indicated that daily contact with environmental water bodies was almost universal across study groups, schistosomiasis infection status may better reflect exposure to high-risk transmission foci than the frequency of water contact alone [16,27]. Differences in the specific water bodies visited, the nature of water contact and the degree of contamination with human and animal excreta are all likely to influence both schistosome transmission and exposure to enteric bacteria. This interpretation is further supported by the finding of increased strain-level microbiome similarity between participants who reported primarily drinking non-piped water compared to those drinking piped water. If the exploratory findings presented here are replicated in future studies, these dynamics may have public health implications during outbreaks of waterborne enteric pathogens such as cholera, where water contamination and flooding caused by natural disasters can facilitate transmission in Malawi [28].

Previous studies investigating the gut microbiota in African schistosomiasis have predominantly relied on 16S amplicon sequencing to characterise taxonomic profiles [2], which is limited by amplification bias, lower taxonomic resolution and reduced sensitivity for detecting low-abundance taxa [29]. Additionally, few studies have evaluated differential abundance using multiple analytical methods to assess the robustness of taxon-specific associations [30]. We used shotgun metagenomic sequencing and multiple analytical methods to support the principal findings, while also incorporating strain-level analyses that are not possible using amplicon sequencing alone. In addition to enabling species- and strain-level analyses, shotgun metagenomics permits profiling of microbial metabolic pathways and ARGs, as well as the recovery of MAGs, providing a more comprehensive view of microbiome structure and function. However, this breadth does not necessarily confer sensitivity comparable to that of targeted culture-based approaches for the detection of low-abundance organisms or ARGs [31]. For instance, no ESBL genes were detected in faecal metagenomic DNA in this study despite the prevalence of gut colonisation with ESBL-producing *E. coli* in Samama being approximately 50% among participants at the time of sampling, as determined by culture-based methods [15]. Consequently, other low-abundance taxa and ARGs may also have remained undetected.

The analyses presented in this study are exploratory in nature and have several limitations. The sample size was small, which in turn limited power for determining subtle shifts in taxonomic, metabolic pathway and resistome profiles. Recruitment from a single endemic community may also limit the generalisability of findings. Furthermore, numerous exploratory analyses were conducted, meaning that some findings may represent false positives despite correcting for multiple testing effects and performing sensitivity analyses. Therefore, validation of results in independent cohorts will be important to confirm these findings and establish their generalisability. Additionally, variables such as diet and social interactions, which are known to influence gut microbiome composition and strain-sharing, respectively, were not measured so could not be accounted for [32,33]. Nevertheless, cases and controls were matched between different households for differential abundance analyses to account for age, sex and co-infection with the common protozoan parasite *G. duodenalis*. The cross-sectional study design also precluded inference of causality or directionality for the association between zoonotic schistosomiasis and *Brachyspira* enrichment. Finally, the primary strain-level analyses were based on pairwise comparisons, in which participants contributed to multiple observations. Although pairwise comparisons are commonly used in microbiome strain-sharing studies [33–35], the resulting pseudoreplication violates the statistical assumption of independence [36]. However, we attempted to mitigate this limitation by confirming significant pairwise associations using participant-level sensitivity analyses.

In conclusion, this study characterises the gut microbiota of children with a newly described form of zoonotic schistosomiasis. In doing so, we identify a potential association with *Brachyspira*, a genus of potential enteric pathogens capable of causing intestinal spirochaetosis. The methods used here also establish a framework for extending schistosome–host microbiome research beyond traditional taxonomic profiling to support inference of strain-level microbiome similarity. Findings demonstrate that strain-level microbiome analyses can reveal patterns consistent with shared environmental exposure that are not apparent from conventional community-level analyses, highlighting their potential to inform integrated WASH interventions aimed at simultaneously reducing transmission of both parasitic and bacterial enteric pathogens in endemic communities.

## Methods

### Ethics statement

Ethical approvals were granted by The College of Medicine Research Ethics Committee, Malawi (approval nos. P.08/21/3381 and P.06/23/4135) and the Liverpool School of Tropical Medicine Research Ethics Committee, United Kingdom (approval no. 22-028). Written informed consent was obtained from the parents or guardians of participants, with additional written assent obtained from participants aged seven years or older.

### Sample and data collection

The samples and data included in this case-control microbiome study were collected within the Hybridization in Urogenital Schistosomiasis (HUGS) longitudinal cohort study that took place between 2021 and 2025. The HUGS investigation was conducted among people aged two years and older in two sentinel, disease-endemic rural villages in southern Malawi (Samama village, Mangochi District [S 14.418767°, E 35.220985°] and Mthawira village, Nsanje District [S 16.849802°, E 35.290041°]). Participants’ household locations were recorded upon initial recruitment to the HUGS study using GPS devices. The HUGS study’s primary objectives were achieved via diagnostic procedures performed using urine samples. However, cross-sectional faecal sampling among SAC (aged 6–15 years) participating in the HUGS study in Samama was performed during the final annual follow-up surveys in June–July 2024.

All 524 SAC retained for follow-up in the HUGS study in Samama (June–July 2024) were invited to provide a faecal sample alongside a paired urine sample at their study visit. We received faecal samples from 265 SAC, of which 247 later passed internal DNA extraction and amplification control. Therefore, a total of 247 faecal samples provided by SAC underwent molecular analysis, alongside paired urine samples [14]. Body temperature measurements were recorded at the time of sample provision. Participant-reported data on current symptoms of fever, antibiotic use within the previous six months, co-morbidities, primary source of drinking water (piped or borehole/other), frequency of contact with environmental water bodies, including freshwater (rivers and lakes) and stagnant water (ponds, marshes, swamps and rice paddies), and site of defaecation (latrine or other), were collected using proforma-based questionnaire interviews. Participants completed questionnaires with the assistance of a parent or guardian.

### Faecal and urine sample processing

Our methodological approach concerning the detection of intestinal schistosomiasis and urogenital schistosomiasis among SAC in Samama (June-July 2024) was recently described in detail [14]. Briefly, DNA was extracted from faecal samples using the QIAamp PowerFecal Pro DNA Kit (QIAGEN). During DNA extraction, each sample was spiked with a standard volume of a phocid herpes virus-1 (PhHV-1) working solution to act as a DNA extraction and PCR internal positive control. A routinely used duplex diagnostic real-time PCR assay [37] targeting a genus-specific *Schistosoma* spp. internal transcribed spacer-2 (ITS2) ribosomal DNA (rDNA) region and a fragment of the PhHV-1 glycoprotein B gene was then used to identify the presence of *Schistosoma* spp. nuclear DNA (nDNA) within these DNA eluates. Following this, a recently developed high-resolution melt (HRM) real-time PCR assay [38] was used to identify mtDNA of the infecting species in ITS2 real-time PCR-positive cases. In addition, urine-egg microscopy was performed using paired urine samples to diagnose urogenital schistosomiasis and categorise infection intensity: negative, 1–9 ova per 10 ml, 10–49 ova per 10 ml and ≥50 ova per 10 ml [14]. Following this, DNA was extracted from egg-positive urine filters using the DNeasy Blood and Tissue Kit (QIAGEN). Urine-filter DNA eluates were then screened to confirm the presence of *Schistosoma* DNA and identify the infecting species using the same methods described above for faecal samples.

### Case-control sample selection for metagenomic sequencing

Faecal DNA eluates analysed within the above-described cross-sectional survey of SAC were considered for shotgun metagenomic sequencing to investigate associations between *S. haematobium* group infections and the gut microbiome. Participants reporting fever or with a body temperature >38 °C, self-reported HIV infection, self-reported tuberculosis infection or *S. mansoni* infection according to PCR-based mtDNA detection were excluded. Ahead of case and control sample selection, we also screened for *Giardia duodenalis* infection using a TaqMan probe real-time PCR assay [39] (primer and probe oligonucleotide sequences, reaction mix and PCR conditions are described in *Supplementary Tables 3–5*). This was done because recent surveys demonstrated that the prevalence of giardiasis is approximately 40% among SAC from Samama [40], and giardiasis has been associated with gut microbiome alterations [41]. Other pathogenic protozoa that could impact the gut microbiome, such as *Cryptosporidium* spp. and *Entamoeba histolytica/dispar*, are very rarely detected in children from Mangochi District [40,42], while no soil-transmitted helminth ova were detected in faecal samples using Kato-Katz faecal microscopy during the HUGS pilot study [13], therefore no further tests for enteric parasitic infections were performed prior to case-control matching and sample selection.

The sample selection strategy was pragmatic and was guided by the number of eligible cases of zoonotic schistosomiasis and by logistical considerations (i.e. sequencing budget). Within the cohort of 247 SAC in Samama [14], there were 12 eligible cases of zoonotic schistosomiasis where faecal *Schistosoma* spp. ribosomal ITS DNA was detected by real-time PCR, and both *S. haematobium* and *S. mattheei* mtDNA signatures, but not *S. mansoni*, were detected in the faeces and/or urine by HRM real-time PCR. SAC with zoonotic schistosomiasis (the ‘ZS’ group) were matched to 12 SAC without schistosomiasis (the ‘Control’ group) and to 12 SAC with ‘classic’ urogenital schistosomiasis caused by *S. haematobium*, from whom no atypical or zoonotic species were detected in urine by microscopy or PCR and no *Schistosoma* spp. ITS2 rDNA was detected in faeces (the ‘UGS’ group). Cases were matched to SAC from different households in the same village based on sex, *G. duodenalis* infection status, and age (with the closest age being selected after meeting the first two matching criteria).

The quantity of selected DNA samples was assessed using a Qubit 4 fluorometer (Thermo Fisher Scientific, USA) with the Invitrogen Qubit double stranded DNA broad range kit, according to the manufacturer’s protocol. NanoDrop One spectrophotometry (Thermo Fisher Scientific) was used to measure DNA purity, and the TapeStation System (Agilent Technologies, USA) was used to measure double stranded DNA fragment size and integrity. DNA eluates were shipped to Genewiz/AZENTA (UK) for library preparation and shotgun metagenomic sequencing using the NovaSeq X platform (Illumina) with a 2 × 150 bp paired-end configuration.

### Read processing and profiling

Raw FASTQ sequence files were quality assessed using FastQC (https://github.com/s-andrews/FastQC; v0.11.9). Trimmomatic [43] (v0.39) was used to remove adapter sequences and low-quality bases with a 4 bp sliding window quality cutoff of Q20 and a minimum read length of 50 bp. Human (host) DNA was filtered out by aligning reads to the *Homo sapiens* reference genome GRCh38.p14 using Bowtie2 [44] (v2.2.5).

Species-level bacterial relative abundance profiles were generated using MetaPhlAn4 [45] (v4.2.2), using the Jan25 marker gene database with default parameters. Relative abundances were expressed as proportions ranging from 0 to 1 using total sum-scale (TSS) normalisation. In addition, microbial metabolic pathway abundance profiles were generated with HUMAnN (v3.9), using default parameters with the native bioBakery3 Jan19 ChocoPhlAn pangenome database and corresponding full UniRef90 database [46]. Pathway abundance profiles were subsequently normalised to copies per million and then expressed as TSS-normalised relative abundances ranging from 0 to 1.

To identify ARGs, reads were aligned to reference sequences in the ResFinder database [47] (v2.3.2) using KMA [48] (v1.3.28). Alignments were then filtered to retain only those with a minimum 90% nucleotide identity and 80% template coverage. For all reference sequences with retained alignment hits after filtering, the corresponding Cluster90 was identified, whereby ResFinder database reference sequences were clustered with a 90% nucleotide sequence identity threshold to identify clusters of highly similar ARGs using CD-HIT [49] (v4.8.1). Normalised abundances of ARGs were calculated from read mapping data, expressed as RPKM (Reads Per Kilobase per Million mapped bacterial reads). These profiles were also TSS-normalised to assess the proportion of the resistome accounted for by each feature.

### Metagenome assembly, binning and classification

*De novo* metagenome assembly was performed with MEGAHIT [50] (v1.2.9) using a *k*-mer spectrum of 31, 59, 87 and 115, selected for efficient recovery of microbial genomes from complex metagenomes [51]. Contiguous sequences (contigs) ≥500 bp were retained, and assembly statistics were calculated with QUAST [52] (v5.0.2). Contigs were screened for ARGs against the ResFinder database using ABRicate (https://github.com/tseemann/abricate; v1.0.1), requiring a minimum of 90% nucleotide identity and 80% template coverage to call a hit.

Contigs were indexed using the Burrows Wheeler Aligner (BWA; v0.7.17), before reads were aligned to the indexed contigs with “bwa-mem” [53]. The contigs and resulting sorted binary alignment map (BAM) files were parsed to the “jgi_summarize_bam_contig_depth” script from MetaBAT2 [54] (v2.15), before the resulting depth files were used by MetaBAT2 to bin assembled contigs of ≥2,500 bp to putative MAGs. CheckM [55] (v1.1.2) was used to assess the completeness and contamination of each MAG. As per criteria set by the Genome Standards Consortium, medium-quality MAGs (>50% completeness and <10% contamination) and high-quality MAGs (>90% completeness and <5% contamination) proceeded to further analysis [56]. MAGs underwent taxonomic classification using GTDB-Tk [57] (v2.1.1) against the Genome Taxonomy Database (GTDB; v09-RS220) and were screened for ARGs using ABRicate, as above. High-quality MAGs were dereplicated at 98% ANI to identify subspecies groups [58] using the ‘skani’ algorithm [59] within dRep (v3.6.2) [60], which chose a representative MAG from each group based on CheckM quality metrics and default parameters.

### Diversity statistics

Data wrangling, analysis, and visualisation were performed in the RStudio environment (R v4.5.2), predominantly using the R/Bioconductor *mia* and *miaViz* (v1.18.0) packages and the R/CRAN *ggplot2* (v4.0.3) and *pheatmap* (v1.0.13) packages. Alpha diversity statistics (Shannon Diversity Index and Gini–Simpson Index) were calculated at the species level. To investigate beta diversity, Bray–Curtis dissimilarity matrices containing pairwise distances among samples were generated from taxonomic relative abundance data and metabolic pathway relative abundance data.

Principal coordinate analysis (PCoA) at the species-level and for metabolic pathways was performed using multi-dimensional scaling to ordinate Bray–Curtis dissimilarity matrices in a reduced-dimensional space. Resulting ordinations were visualised with 95% t-distribution ellipses stratified by schistosomiasis category. To investigate associations of schistosomiasis category, giardiasis, age group (6–10 and 11–15), sex, antibiotic use within the previous six months and primary drinking water source with both species- and genus-level bacterial taxonomic profiles and metabolic pathways, Permutational Multivariate Analysis of Variance (PERMANOVA) and homogeneity tests were performed upon Bray-Curtis dissimilarity matrices using the *mia* ‘getPERMANOVA’ function, with 9,999 permutations. Statistical significance was determined at *P* <0.05.

### Differential abundance analysis

For exploratory differential abundance analysis of gut microbiome features comparing between groups, MaAsLin2 was used as the primary method, implemented in the R/Bioconductor *MaAsLin2* package [18] (v1.15.1). Analyses were performed on taxonomic profiles at the genus and phylum levels, as well as metabolic pathway profiles and ARG profiles at the class and Cluster90 levels. Taxonomic and metabolic pathway profiles were analysed as log_2_-transformed TSS-normalised relative abundance profiles, whereas ARG profiles were analysed as log_2_-transformed RPKM-normalised abundances.

Low-prevalence and low-abundance features were removed before model fitting whereas features with a relative abundance of >0.1% in >25% of samples were retained. In mixed-effects models, schistosomiasis category was included as a fixed effect, while ‘MatchID’ was included as a random intercept to account for the matched study design (therefore each sample was compared to a matched sample from a different household).

As a sensitivity analysis, differential abundance analysis of taxonomic and metabolic pathway relative abundance data was repeated using LinDA [19] in the R/Bioconductor *MicrobiomeStat* (v1.4) package. For both MaAsLin2 and LinDA models, the Benjamini–Hochberg procedure was used to control the false discovery rate (FDR). Associations with FDR-adjusted *P*-values (*q*-values) <0.10 were considered candidate features for further investigation, whereas associations with *q*-values <0.05 were considered statistically significant. Concordance in the direction and significance of associations between MaAsLin2 and LinDA was used to assess the robustness of the findings.

For features identified by differential abundance analyses as associated with zoonotic schistosomiasis, Spearman’s rank tests were used to assess correlations between feature relative abundance and normalised faecal *Schistosoma* DNA abundance. To derive this measure, *Schistosoma* DNA abundance was normalised to the PhHV-1 internal control using the ‘2^-ΔCt^’ method: ΔCt was calculated as the cycle threshold (Ct) value for the *Schistosoma* spp. ITS2 target minus the Ct value for the PhHV-1 glycoprotein B target. This approach accounts for variation in DNA recovery and PCR inhibition between samples and transforms Ct differences to a linear measure proportional to relative *Schistosoma* DNA abundance [61].

### Strain-level microbiome similarity profiling

Strain-level profiling was performed using StrainPhlAn scripts within the MetaPhlAn environment [45] for SGBs detected in >20% of study samples. Dominant-strain consensus marker sequences were reconstructed and used to generate SGB-specific maximum-likelihood phylogenies (--phylophlan_mode accurate), retaining samples meeting marker coverage criteria (≥25% of markers and ≥10 markers after filtering). To identify highly similar dominant strains shared by participants in our study, phylogenies incorporated faecal metagenomes from people not receiving antibiotics (n = 53) in an external reference gut metagenomic study carried out in Blantyre, Malawi [62]. This population, separated geographically (∼150 km) and temporally (>5 years), provided background genetic distances between epidemiologically unrelated strains. Using the “strain_transmission.py” script, SGB-specific similarity thresholds were defined using the lower tail normalised phylogenetic distances between unrelated strains (default: 3rd percentile). Pairs of samples in our study with phylogenetic distances below the corresponding SGB threshold were classified as harbouring highly similar dominant strains (*Supplementary Figure 4*).

Strain-level microbiome similarity was calculated for each participant pair as the percentage of SGBs profiled in both samples in which dominant strains were deemed highly similar by StrainPhlAn. Strain-level microbiome similarity was calculated for all 630 pairwise comparisons among the 36 study participants. Three within-household pairs were then excluded to minimise potential confounding from increased strain sharing among household members [33,34], leaving 627 pairs for downstream analyses.

### Statistical analysis of strain-level microbiome similarity

Bivariate associations with strain-level microbiome similarity were assessed using Spearman’s rank correlation for geographic distance between households and Kruskal–Wallis tests with post-hoc Dunn’s tests (Benjamini–Hochberg correction) for categorical variables. These included schistosomiasis status (both infected, neither infected, discordant), schistosomiasis category among infected participants (both UGS, both ZS, discordant), giardiasis status (both infected, neither infected, discordant), age difference (<2, 2–3, 4–5, 6–7, or 8–9 years), sex composition (both male, both female, discordant), and piped water as the primary drinking water source (both yes, both no, discordant). To assess whether significant pairwise associations persisted at the participant level, sensitivity analyses were performed for variables with significant Kruskal–Wallis tests (*P* <0.05). For each participant, median similarity with participants in the same category (within-group) and different categories (between-group) was calculated, and their difference compared between groups using Wilcoxon rank-sum tests.

The number of highly similar dominant strains was modelled as the outcome, with the logarithm of the number of jointly profiled SGBs included as an offset. Schistosomiasis status, sex composition, age difference and geographic distance were included *a priori*, alongside additional covariates significant in both pairwise bivariate and participant-level sensitivity analyses. Exponentiated coefficients were reported as IRRs with 95% confidence intervals. Permutation-based variable importance was assessed by independently permuting each covariate across participant pairs 1,000 times, refitting the model while retaining all other covariates unchanged. Importance was quantified as the reduction in model log-likelihood relative to the original model, with larger positive values indicating greater contributions to model fit.

### Lower bound SNP distance estimation

TRACS [20] (v1.0.1) was used to estimate genetic divergence between strains of selected species of interest (i.e., *Brachyspira* spp.) based on single nucleotide polymorphism (SNP) distances, using high-quality study-derived MAGs as references. Unlike StrainPhlAn, which relies on a dominant consensus strain, TRACS uses whole-genome alignments and accounts for within-host strain diversity, enabling the estimation of conservative lower bound SNP distances when multiple strains of the same species are present within a sample. SNP distance estimates were retained for downstream analysis only if they were calculated from informative sites representing >50% of the respective reference genome.

## Author statements

### Data availability

Sequencing data generated from all samples in this study have been deposited to National Center for Biotechnology Information (NCBI) within the Sequence Read Archive (SRA) under BioProject PRJNA1532595. Data and code for analyses carried out in R are available at: https://github.com/amoreo71/malawi_schisto_microbiome.

### Author contributions

Conceptualisation: AMO’F, JM, JRS, APR; Methodology: AMO’F, JRS, APR; Data curation: AMO’F, DL, PM, GN, JML, SAK, JM, JRS; Formal analysis: AMO’F, DL, PM, GN, LJC, JA, JRS, APR; Resources: AB, SCW, JM, JRS, APR; Writing – original draft: AMO’F; Writing – reviewing and editing: all authors.

### Competing interests

None to declare.

### Funding

AMO’F is funded by the Medical Research Council via the Liverpool School of Tropical Medicine / Lancaster University doctoral training partnership (grant no. MR/W007037/1). The HUGS study was funded by the Wellcome Trust within a Joint Investigator Award (grant no. 220818/Z/20/Z). APR acknowledges funding from the Medical Research Council, Biotechnology and Biological Sciences Research Council and Natural Environment Research Council which are all Councils of UK Research and Innovation (grant no. MR/W030578/1) under the umbrella of the JPIAMR (Joint Programming Initiative on Antimicrobial Resistance), and UKRI through the Strength in Places Fund (grant no. SIPF 36348). The funders had no role in study design, data collection and analysis, decision to publish or preparation of the manuscript.

## Acknowledgements

We are grateful to all participants in the study. We thank staff associated with the HUGS study from Liverpool School of Tropical Medicine and Malawi Liverpool Wellcome Research Programme, including Alexandra Juhász, Sam Jones, Ruth Cowlishaw, Sarah Rollason, Priscilla Chammudzi, Donales R. Kapira, E. James LaCourse, Bright Mainga and Clinton Nkolokosa.

## Notes

https://github.com/amoreo71/malawi_schisto_microbiome

### Competing Interest Statement

The authors have declared no competing interest.

