## Supplementary Information for "Strain-resolved metagenomics shows gut microbiome signatures in a case-control study of Malawian children with zoonotic schistosomiasis"

**Supplementary Table 1. Summary of Illumina sequencing data and metagenome assemblies.**
The number of bacterial read pairs is an estimate generated by MetaPhlAn.

| ID | Illumina short reads | | | Assembly (contigs ≥500 bp) | | | Metagenome-assembled genomes | |
|---|---|---|---|---|---|---|---|---|
| | Total read pairs | Read pairs passing QC | Bacterial read pairs passing QC | Contigs | Total length | N50 | High-quality | Medium-quality |
| C1 | 45856548 | 42744313 | 33753902 | 358103 | 592797817 | 2436 | 23 | 42 |
| C2 | 51090686 | 47745199 | 38263771 | 204734 | 382379999 | 3369 | 9 | 36 |
| C3 | 52134191 | 48990125 | 40133416 | 283446 | 493050278 | 2806 | 15 | 37 |
| C4 | 49720304 | 46380960 | 38154358 | 227667 | 406575794 | 3046 | 11 | 44 |
| C5 | 46384051 | 43250309 | 34361676 | 320964 | 530539627 | 2423 | 14 | 42 |
| C6 | 49820859 | 46725988 | 37678605 | 330217 | 551908726 | 2558 | 17 | 52 |
| C7 | 49400966 | 45902484 | 35252640 | 325828 | 555785021 | 2593 | 18 | 40 |
| C8 | 50446606 | 46591642 | 37953642 | 368635 | 661465016 | 3001 | 31 | 57 |
| C9 | 45033192 | 42058384 | 34229975 | 265055 | 514185073 | 3454 | 22 | 46 |
| C10 | 54060654 | 50637552 | 41121583 | 280767 | 475554875 | 2606 | 11 | 34 |
| C11 | 52727129 | 49483784 | 37204116 | 377414 | 654829394 | 2747 | 18 | 59 |
| C12 | 51692354 | 48388320 | 36213037 | 227529 | 448442071 | 3756 | 17 | 40 |
| U1 | 46741922 | 43796259 | 34773156 | 111121 | 250625387 | 5426 | 14 | 22 |
| U2 | 48829099 | 45992845 | 34927090 | 178667 | 327931669 | 3314 | 8 | 31 |
| U3 | 51227282 | 47795731 | 36751460 | 460950 | 709141912 | 2052 | 16 | 51 |
| U4 | 55659237 | 51946951 | 40831366 | 413473 | 745583976 | 3124 | 37 | 61 |
| U5 | 46505354 | 42857200 | 35319544 | 283766 | 536113609 | 3419 | 24 | 44 |
| U6 | 50663427 | 47131949 | 35918702 | 430363 | 654493226 | 2048 | 24 | 28 |
| U7 | 41125582 | 37982515 | 31450347 | 299322 | 553466786 | 3348 | 33 | 58 |
| U8 | 52227745 | 48895036 | 37179837 | 442484 | 732140838 | 2442 | 22 | 55 |
| U9 | 50393030 | 47020023 | 36452889 | 377120 | 641549029 | 2625 | 24 | 52 |
| U10 | 46335287 | 43430372 | 34183432 | 149126 | 357451828 | 5991 | 15 | 29 |
| U11 | 52894325 | 49770742 | 39586393 | 284728 | 478298797 | 2596 | 12 | 41 |
| U12 | 53621088 | 49713363 | 40509692 | 334946 | 585365571 | 2699 | 17 | 50 |
| Z1 | 54359282 | 51575420 | 41327186 | 305478 | 586030693 | 3417 | 33 | 44 |
| Z2 | 52531956 | 49857048 | 39877111 | 326951 | 551967777 | 2498 | 18 | 45 |
| Z3 | 51037726 | 48221799 | 38573584 | 340325 | 552861644 | 2386 | 18 | 48 |
| Z4 | 52343077 | 49379109 | 39433825 | 374688 | 620607393 | 2450 | 29 | 52 |
| Z5 | 50900339 | 48424468 | 37978607 | 289493 | 516933645 | 3015 | 18 | 46 |
| Z6 | 52291098 | 49773985 | 39164277 | 391004 | 648864146 | 2444 | 20 | 60 |
| Z7 | 51583734 | 48656337 | 38615121 | 329340 | 653330736 | 3899 | 27 | 56 |
| Z8 | 53500340 | 50834035 | 39510437 | 245008 | 470305987 | 3619 | 19 | 46 |
| Z9 | 56287467 | 53353768 | 40038105 | 278939 | 455592213 | 2371 | 6 | 31 |
| Z10 | 55229122 | 52666800 | 38948948 | 300066 | 503453817 | 2583 | 15 | 50 |
| Z11 | 59224576 | 55993868 | 45667404 | 264871 | 452741685 | 2622 | 20 | 29 |
| Z12 | 48940355 | 46190121 | 37045500 | 360360 | 598055828 | 2526 | 26 | 42 |

**Supplementary Table 2. Multivariable negative binomial regression model outputs for factors associated with strain-level microbiome similarity.**

The table presents regression coefficient estimates, standard errors, and *P*-values for associations between participant pair characteristics and the number of highly similar dominant strains shared. These model outputs were used to calculate incident rate ratios (IRRs) and 95% confidence intervals, as shown in Figure 5 of the main manuscript. The model included the logarithm of the total number of shared species-level genome bins (SGBs) as an offset. Age difference is in years. Household distance was modelled per 100 m increase.

| **Term** | **Estimate** | **Standard error** | ***P*-value** |
|---|---|---|---|
| Intercept | -2.368 | 0.102 | <0.001 |
| category_contrast_Control×UGS | 0.114 | 0.078 | 0.142 |
| category_contrast_Control×ZS | 0.065 | 0.078 | 0.402 |
| category_contrast_UGS×UGS | 0.250 | 0.091 | 0.006 |
| category_contrast_UGS×ZS | 0.264 | 0.077 | <0.001 |
| category_contrast_ZS×ZS | 0.218 | 0.090 | 0.015 |
| age_diff_2-3 | -0.006 | 0.054 | 0.905 |
| age_diff_4-5 | -0.038 | 0.060 | 0.520 |
| age_diff_6-7 | -0.010 | 0.060 | 0.866 |
| age_diff_8-9 | -0.076 | 0.122 | 0.532 |
| sex_contrast_Discordant | 0.091 | 0.062 | 0.139 |
| sex_contrast_Both male | 0.184 | 0.064 | 0.004 |
| piped_water_contrast_Discordant | -0.085 | 0.047 | 0.068 |
| piped_water_contrast_Both piped | -0.130 | 0.062 | 0.038 |
| distance_100m | -0.033 | 0.013 | 0.014 |

**Supplementary Table 3. Primer and probe sequences used to amplify a fragment of the *Giardia duodenalis* 18S ribosomal RNA gene and a fragment of the PhHV-1 glycoprotein B gene.**

| Primer/probe | Target (fragment size) | Oligonucleotide sequence (5' – 3') |
|---|---|---|
| Giardia-80_F * | *G. duodenalis* 18S (62 bp) | GACGGCTCAGGACAACGGTT |
| Giardia-127_R † | | TTGCCAGCGGTGTCCG |
| Giardia-105_Pro ‡ | | (FAM¥) CCCGCGGCGGTCCCTGCTAG |
| PhHV-1-267s_F * | PhHV-1 glycoprotein B (89 bp) | GGGCGAATCACAGATTGAATC |
| PhHV-1-337as_R † | | GCGGTTCCAAACGTACCAA |
| PhHV-1-305tq_Pro ‡ | | (Cy5¥) TTTTTATGTGTCCGCCACCATCTGGATC |

*Forward primer, †Reverse primer, ‡*Probe,* ¥*Fluorophore*

**Supplementary Table 4. Reaction mix used to carry out duplex TaqMan probe real-time PCR targeting a fragment of the *Giardia duodenalis* 18S ribosomal RNA gene and a fragment of the PhHV-1 glycoprotein B gene.**

| Reagent | Concentration (reaction mix) | Volume per sample (µl) |
|---|---|---|
| $ddH_2O$ | / | 2.92 |
| Giardia-80_F * (10 µM) | 0.25 µM | 0.30 |
| Giardia-127_R † (10 µM) | 0.25 µM | 0.30 |
| Giardia-105_Pro ‡ (10 µM) | 0.1 µM | 0.12 |
| PhHV-1-267s_F * (10 µM) | 0.1 µM | 0.12 |
| PhHV-1-337as_R † (10 µM) | 0.1 µM | 0.12 |
| PhHV-1-305tq_Pro ‡ (10 µM) | 0.1 µM | 0.12 |
| AppPROBE No ROX 2X Mix (Appleton Woods) | / | 6 |
| DNA | / | 2 |
| TOTAL | / | 12 |

*Forward primer, †Reverse primer, ‡*Probe,* ¥*Fluorophore*

**Supplementary Table 5. Cycling conditions used to carry out duplex TaqMan probe real-time PCR targeting a fragment of the *Giardia duodenalis* 18S ribosomal RNA gene and a fragment of the PhHV-1 glycoprotein B.**

| Cycles | Time | Temperature |
|---|---|---|
| 1 | 3 min | 95 °C |
| 40 | 10 sec | 95 °C |
| | 30 sec | 60 °C |

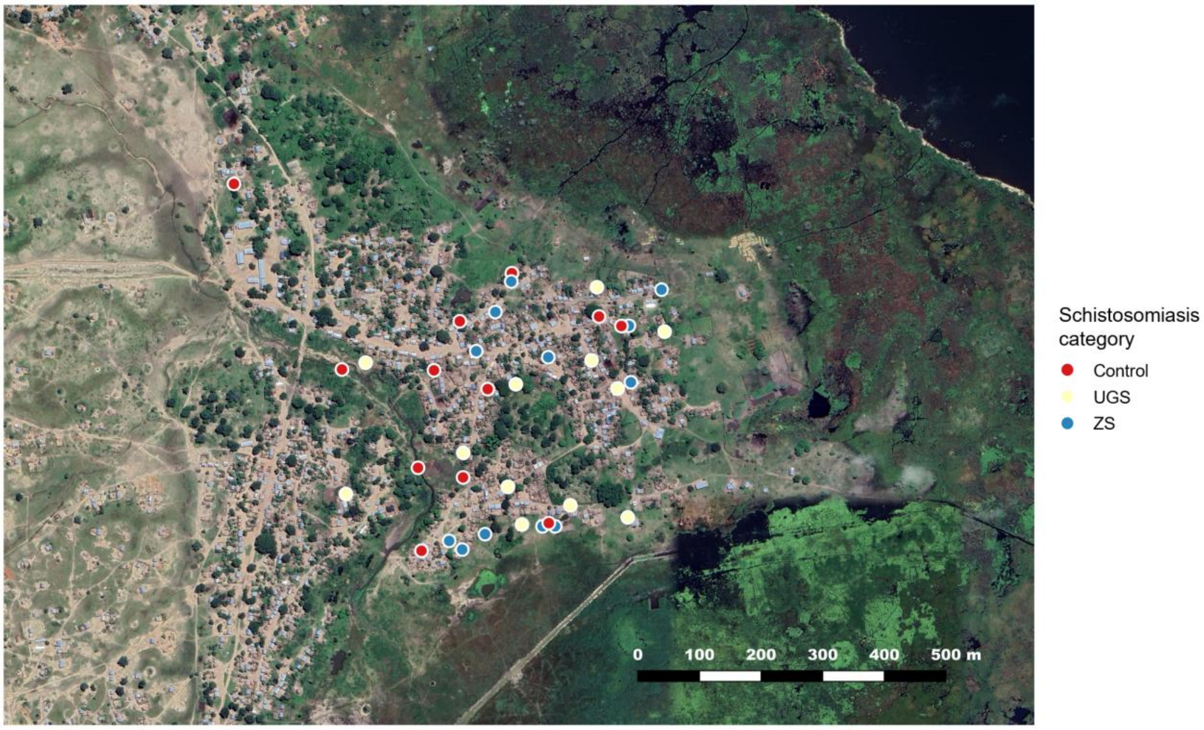

**Supplementary Figure 1. Approximate household locations of study participants in Samama, Mangochi District, Malawi.**

Household locations of the 36 SAC included in this metagenomic study are coloured by participant schistosomiasis category: Control (no schistosomiasis), ZS (zoonotic schistosomiasis) and UGS (urogenital schistosomiasis). To protect participant confidentiality, household GPS coordinates were anonymised by applying a small random offset (±50 m) to both latitude and longitude prior to plotting. Satellite imagery was obtained from Google (©2026 Airbus, CNES / Airbus, Maxar Technologies).

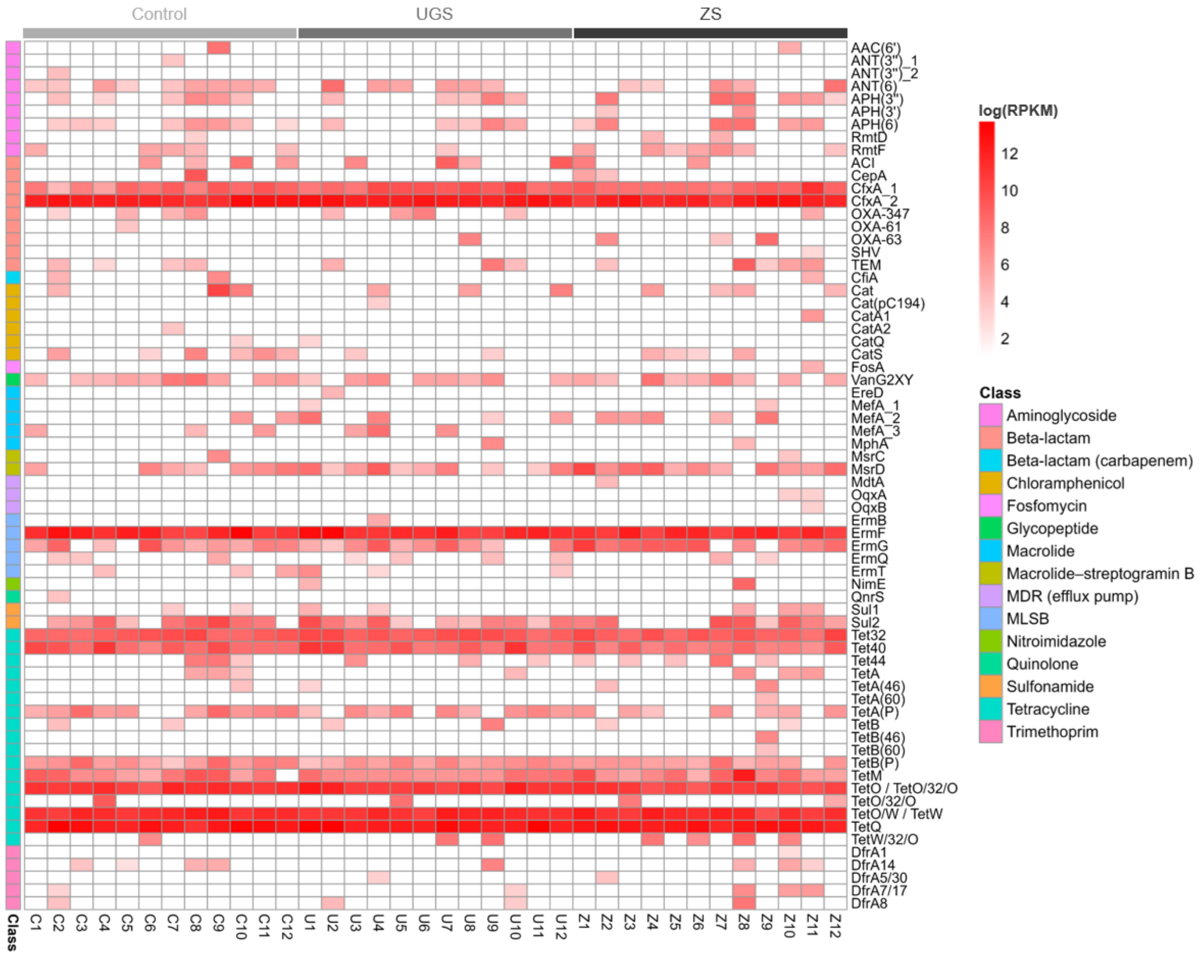

**Supplementary Figure 2. Heatmap of normalised ARG Cluster90 abundances.**
Normalised ARG abundances are shown as log-transformed RPKM. In row annotations, ARG Cluster90s are colour-coded by the antibiotic class to which they are predicted to confer resistance. Samples from the 'Control', 'UGS' and 'ZS' groups begin with 'C', 'U' and 'Z', respectively. MDR = multidrug-resistant.

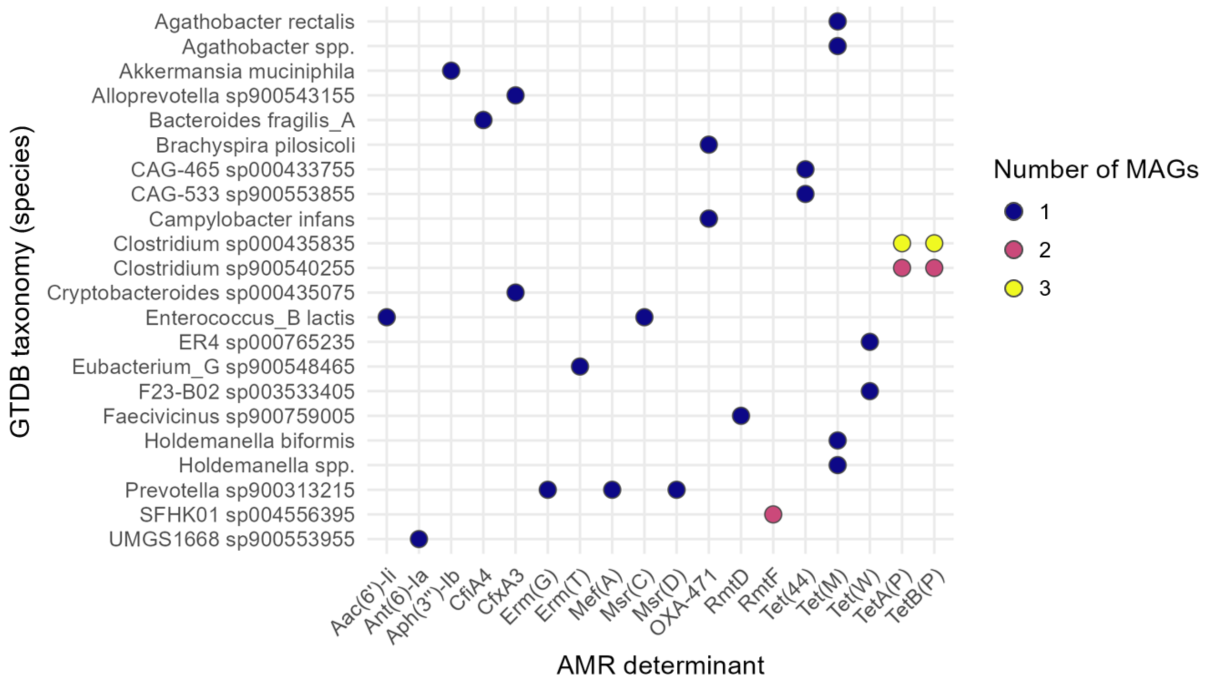

**Supplementary Figure 3. ARGs identified in high- or medium-quality MAGs.**
The colour scale indicates the number of MAGs the ARG was identified in.

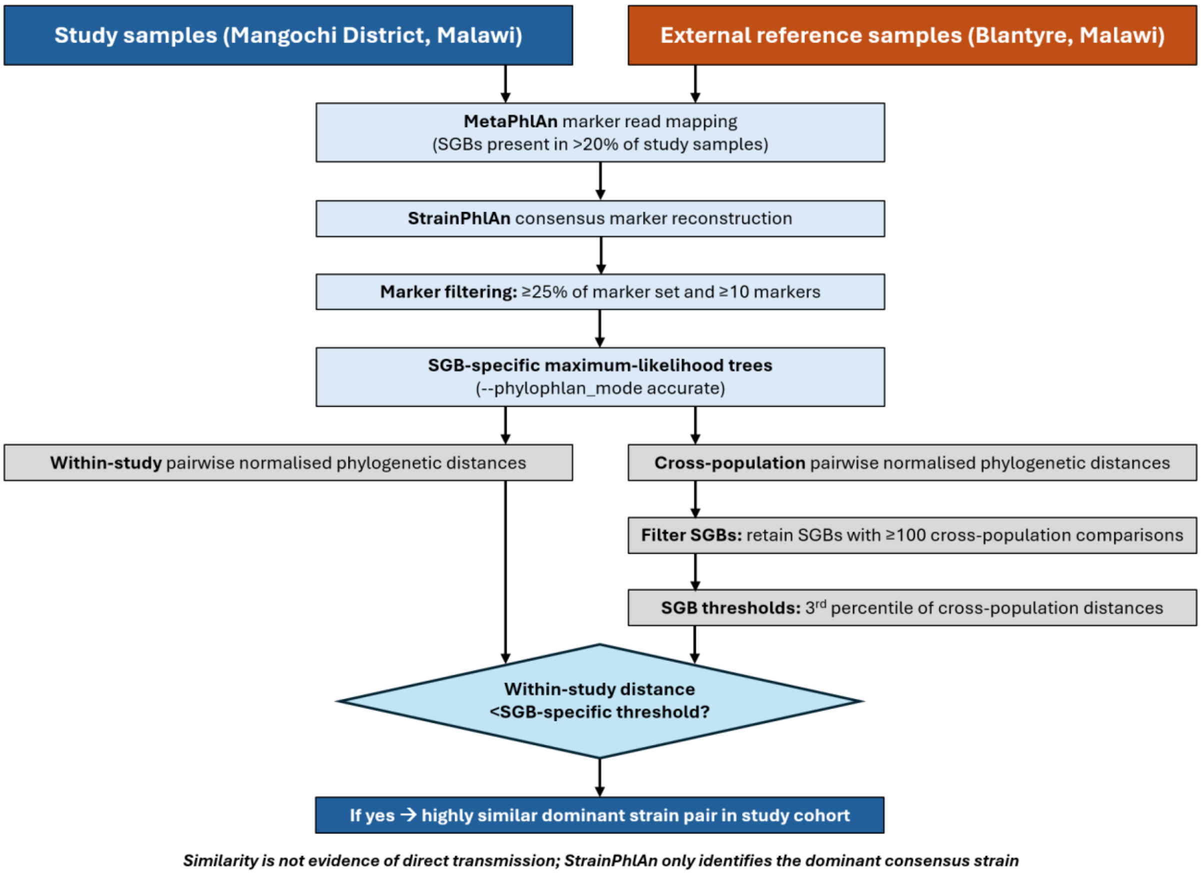

**Supplementary Figure 4. Identification of highly similar dominant strains in the gut microbiomes of study participants.**
External reference samples from Blantyre, Malawi (n = 53) were used to identify background genetic distances between epidemiologically unrelated strains. External samples were only included from people not receiving antibiotic treatment as antibiotics may cause perturbation of the gut microbiome, including loss or replacement of resident strains. Where multiple external samples provided by the same individual were available, a single representative sample was selected, defined as the first sample passing sequencing quality control (typically the baseline timepoint). Accessions for the external samples are available from: https://github.com/amoreo71/malawi_schisto_microbiome/blob/main/Data/blantyre_external_reference_samples_metadata.csv.
